# Phenotypic and Molecular characterization of thermophlic bacterial isolates from the rizosphere of *Panicum antidotale* in Cholistan desert, Pakistan

**DOI:** 10.1101/2025.10.16.682883

**Authors:** Muhammad Aslam, Misbah Afzal, Rimsha Batool, Tanzila Rehman, Faiz-ul-Hassan Nasim, Samina Ejaz

## Abstract

The major aim of this study was to isolate, characterize, and determine the distribution of culturable bacteria in the rhizosphere of *Panicum antidotale* (PAMA) from the Cholistan desert, Pakistan. A total of 109 thermophilic bacterial isolates were obtained from PAMA and subjected to morphological, biochemical, and molecular characterization at The Islamia University of Bahawalpur. Standard protocols were used for physical and biochemical analysis, while molecular identification involved 16S rRNA profiling, restriction fragment length polymorphism (RFLP), and DNA sequence analysis. PCR amplicons of selected isolates with distinct RFLP patterns were sequenced, and the DNA sequences were analyzed using BLAST to determine species identity. Results revealed the presence of *Bacillus subtilis*, *Brevibacillus borstelensis*, and *Paenibacillus dendritiformis* in the majority of analyzed rhizosphere samples. *Bacillus licheniformis* and *Bacillus subtilis* were the most dominant species in PAMA, accounting for 45% and 30% of the isolates, respectively. In addition, lower frequencies of other species, such as *Paenibacillus dendritiformis* (10–15%), were also observed. These findings demonstrate that *Panicum antidotale* supports a unique rhizospheric bacterial community dominated by thermophilic *Bacillus* species. This study represents the first comprehensive investigation of PAMA microbiota in the Cholistan desert. The results provide novel insights into bacterial diversity in extreme environments and highlight the potential role of these communities in plant growth promotion and stress adaptation. Further studies are warranted to explore the ecological significance and possible biotechnological applications of PAMA-associated thermophilic bacteria for sustainable agriculture and environmental management.

**Importance:** This study presents the first molecular-level investigation of thermophilic bacterial communities inhabiting the rhizosphere of *Panicum antidotale* in the Cholistan Desert, Pakistan. Through combined phenotypic, biochemical, and molecular analyses, diverse thermophilic *Bacillus*, *Brevibacillus*, and *Paenibacillus* species were identified as dominant members of this extreme arid ecosystem. These bacteria possess plant growth-promoting and stress-alleviating traits, underscoring their ecological significance in supporting desert plant resilience. Importantly, this first molecular characterization of *P. antidotale*-associated microbiota reveals that these isolates may also serve as promising sources of valuable enzymes and bioactive metabolites. The findings broaden our understanding of microbial diversity in arid rhizospheres and highlight the potential applications of these thermophilic bacteria in sustainable agriculture, biotechnology, and environmental management.

## Introduction

Plant roots interact closely with a distinct community of soil microbes residing in the root-adjacent zone called the rhizosphere. This region ranks among Earth’s most intricate ecosystems, hosting millions of microbial cells whose populations fluctuate based on the plant’s genetic makeup and developmental stage. Within the rhizosphere, roots release diverse chemical compounds that attract soil microorganisms. The rhizobacteria living there help plants withstand environmental stresses (abiotic stresses) through multiple mechanisms. These include modifying plant hormone levels, inducing metabolic changes, boosting antioxidant defenses, producing bacterial exopolysaccharides (EPS), and protecting and enhancing root development. Furthermore, these microbes can influence the production of plant metabolites, enhance photosynthesis, and increase carbohydrate and protein content, ultimately improving plant yield traits even under stressful conditions [1]. Plant growth-promoting rhizobacteria (PGPR) are free-living bacteria that colonize plant roots and promote plant growth. PGPR may promote plant growth by using their own metabolism (solubilizing phosphates, producing hormones, or fixing nitrogen), by directly affecting the plant metabolism (increasing the uptake of water and minerals), enhancing root development, increasing the enzymatic activity of the plant, by “helping” other beneficial microorganisms to enhance their action on the plant, or by suppressing plant pathogens [2]. Numerous bacterial genera including *Bacillus, Azotobacter, Pseudomonas, Alcaligenes, Arthobacter, Klebsilla, Azospirillum, Enterobacter,* and *Rhizobium*, have been documented for their ability to promote plant growth [3]. *Panicum antidotale* is a dominant, highly prevalent, and nutritious grass species exhibiting broad ecological tolerance. Consequently, it thrives in both typical conditions and drought-affected landscapes. Populations of *P. antidotale* found in arid zones of the Cholistan desert or on non-stressed soils have likely endured varying levels of water scarcity over extended periods [4]. This plant develops a robust rhizome-based root system, particularly after multiple harvests, enhancing its drought tolerance. Panicgrass (*Panicum antidotale*) is highly salt-resistant and can produce significant yields even under highly saline conditions [5]. It contains several bioactive compounds, including flavonoids and phenols. These phytochemicals are associated with traditional medicinal uses such as managing diabetes, reducing anxiety, relieving pain, fighting bacteria, acting as antioxidants, and reducing inflammation. It has also been applied topically as a poultice to soothe localized inflammation and painful conditions like joint pain [6]. It is a medicinal plant and due to its distinct biological features, the plant is anticipated to harbor biotechnologically significant microorganisms within its root zone. 16S ribosomal RNA gene sequencing analyzes specific genetic regions to classify bacteria/archaea taxonomically and estimate microbiome diversity. This method examines only a small portion of microbial DNA yet reveals valuable insights into community composition and identification. The 16S rRNA gene encodes part of the small (30S) subunit of prokaryotic ribosomes (which consist of 50S and 30S subunits). As a universally conserved gene in bacteria and archaea, it acts as a “molecular clock” due to its stable function in cells and variable regions that enable phylogenetic analysis and tracking of evolutionary divergence [7]. The choice of PCR primers significantly influences observed microbial diversity in a study. To amplify a target region of the 16S rRNA gene, “broad-range” (universal) primers bind to conserved sequences flanking the desired hypervariable zone [8].

The rhizosphere microbiome of the Cholistan desert remains largely unexplored, representing a critical gap in understanding microbial ecology within extreme arid environments. This pioneering study focuses on *Panicum antidotale*, a resilient desert grass, to characterize its culturable rhizobacterial community. We aim to identify unique bacterial species endemic to the Cholistan, map their distribution across native grasses, decipher plant-bacteria interactions, and illuminate the overall diversity sustaining these desert ecosystems. Our findings will provide foundational insights into microbial adaptations supporting plant survival in hyper-arid conditions, revealing potential novel extremophiles and their ecological roles.

## Material and Methods

### Study Area

The research was conducted in the Cholistan Desert, locally known as Rohi, which encompasses 2.6 million hectares. Stretching 480 km southeast toward the Nara Desert in Sindh, its width ranges from 32 to 192 km within the coordinates 27°42’N to 29°45’N and 69°52’E to 75°24’E. More than four-fifths (81%) of the terrain consists of vast sand dunes, while the remaining 19% comprises alluvial flats and sand hummocks. The desert is administratively divided into Greater and Lesser Cholistan, covering parts of Bahawalpur, Rahim Yar Khan, and Bahawalnagar districts. Hardy xerophytic plants thrive across diverse soil and climatic conditions, with more notable growth in the arid east compared to the hyperarid southern region [9].

### Rhizospheric Soil Collection

To isolate bacteria, 10 g of rhizospheric soil was collected from *Panicum antidotale* grass in the Cholistan Desert during three seasons: extreme summer (June–July), winter (December– January), and the rainy season. Samples were collected using sterile techniques (ethanol-flamed spatulas, sterile plastic bags), stored immediately at 4°C in a SANYO MPR-311D(H) incubator, and processed within seven days. Each sample was preserved in labeled sterile jars at 4°C for further investigation of culturable bacterial populations.

### Cultivable Bacterial Growth

Bacterial isolation and quantification began with the preparation of a microbial suspension by adding a measured amount of rhizospheric soil to sterile Ringer’s solution in a cotton-plugged flask. This suspension was homogenized by continuous swirling to ensure uniformity. To estimate viable populations, serial dilutions (10⁻¹–10⁻⁶) were prepared, as microscopic counts cannot distinguish living from dead cells. A 100 µl aliquot of appropriate dilutions (e.g., 10⁻³– 10⁻⁶) was spread on LB agar plates, the standard medium used for this purpose. Plates were incubated at 30°C for 16–24 hours for mesophilic bacteria or at 50°C for thermophiles. After incubation, visible colonies with varying colors and morphologies were observed; clear zones around some colonies indicated competitive inhibition. Colonies were counted, and the average number was used to calculate Colony Forming Units per gram of soil (CFU/g) using the formula:

CFU/g soil = Number of colonies × Dilution Factor

Distinct colonies were isolated as pure strains and stored at –70°C in glycerol for future study.

### Strain Purification

Distinct colonies from LB agar plates were selected and repeatedly streaked on fresh plates until pure isolates were obtained. These purified bacterial cultures were preserved long-term as glycerol stocks (50% LB medium, 50% glycerol) at –70°C in a PLATINUM-550 freezer. Overnight LB broth cultures were prepared from these stocks for further analysis. The isolates were characterized using standard microbiological, biochemical, and molecular techniques.

### Biochemical Characterization of Bacterial Isolates

Isolates were characterized microscopically and biochemically. Microscopic analysis included colony morphology, Gram staining, motility, and cell shape. Biochemical tests assessed the production of enzymes and metabolites including amylase, catalase, protease, gelatinase, urease, indole, hydrogen sulfide (H₂S), and antibiotics. Additional tests included carbohydrate fermentation, the Ulrich Milk Test (to evaluate bacterial transformation of milk), and MR-VP broth tests.

### Genomic DNA Extraction

L.B. broth cultures of purified isolates were processed to extract genomic DNA using a standard protocol consisting of four stages: disruption, lysis, removal of proteins/contaminants, and DNA recovery. The extracted DNA was used for amplification of the 16S rRNA gene via PCR. SDS-based extraction was performed, and DNA integrity was checked by agarose gel electrophoresis (0.8–1%) using TAE buffer (pH 8). DNA was quantified spectrophotometrically (Model-APEL-PD303UV) by measuring absorbance at 260 nm.

### Amplification of the 16S rRNA Gene

In vitro amplification was performed using a thermocycler (MY GENETm Model MG-96+). PCR conditions were optimized for 16S rRNA gene amplification using universal primers reported in earlier studies [10]:

- Sense primer: 5’-AACACATGCAAGTGGAAC-3’ (positions 50–67)
- Antisense primer: 5’-ACGGGCGGTGTGTACAAG-3’ (positions 1406–1389)

The expected amplicon size using *E. coli* DNA was 1357 bp. Extracted DNA was diluted to 20 ng/µl with sterile water. Each 50 μl reaction contained: 1× PCR buffer, 1.5 mM MgCl₂, 100 μM dNTP mix, 0.3 μl of each primer (10 pmol), 1.25 U of Taq DNA polymerase, and the appropriate amount of chromosomal DNA. The PCR program included:

- Initial denaturation: 5 min at 94°C
- 40 cycles of: 30 s at 94°C (denaturation), 60 s at 50°C (annealing), 60 s at 72°C (extension)
- Final extension: 10 min at 72°C

### Amplicon Analysis via RFLP

RFLP analysis of 16S rRNA PCR amplicons was performed using 4-base cutter restriction enzymes: RsaI, HpaII, TaqI, HinfI, and HhaI. Gel images were acquired using a UVpro gel documentation system. Restriction fragments were analyzed with PAST-3 software to generate dendrograms. Gels were coded with 1 and 0 to indicate the presence or absence of bands, and data were transferred to an Excel sheet for dendrogram construction.

### Phylogenetic Analysis of 16S rRNA

PCR products of representative isolates were selected for sequencing based on dendrogram analysis. Two to three isolates from each group were sent to Macrogen, South Korea, for sequencing. The sequences obtained were analyzed via BLAST [11]. Alongside the 20 sequences generated, 12 closely related sequences were retrieved from GenBank. A total of 33 sequences were aligned using ClustalX and refined manually in BioEdit. Phylogenetic trees were constructed with MEGA7 [12] using the Neighbor-Joining method [13], and reliability was assessed with 100 bootstrap replicates. Evolutionary distances were calculated using the p-distance method. Positions with gaps or missing data were removed, resulting in 682 positions for the final analysis [14].

## Results

### Phenotypic characterization

Morphological analysis revealed both consistent and variable traits among the bacterial isolates (Table 1). All isolates exhibited non-chromogenic pigmentation (no pigment production) and were Gram-negative. The predominant cell shape was bacillus, accounting for 88% of isolates, while cocci represented 12%. No spirilla were observed. Colony color was primarily off-white (60%), with the remaining 40% being non-off-white. Colony margins showed diversity, with 39% being branching and 61% non-branching. Colony elevation was divided between raised (39%) and flat (61%), while opacity was classified as opaque (36%) or non-opaque (64%).

**Table 1.**
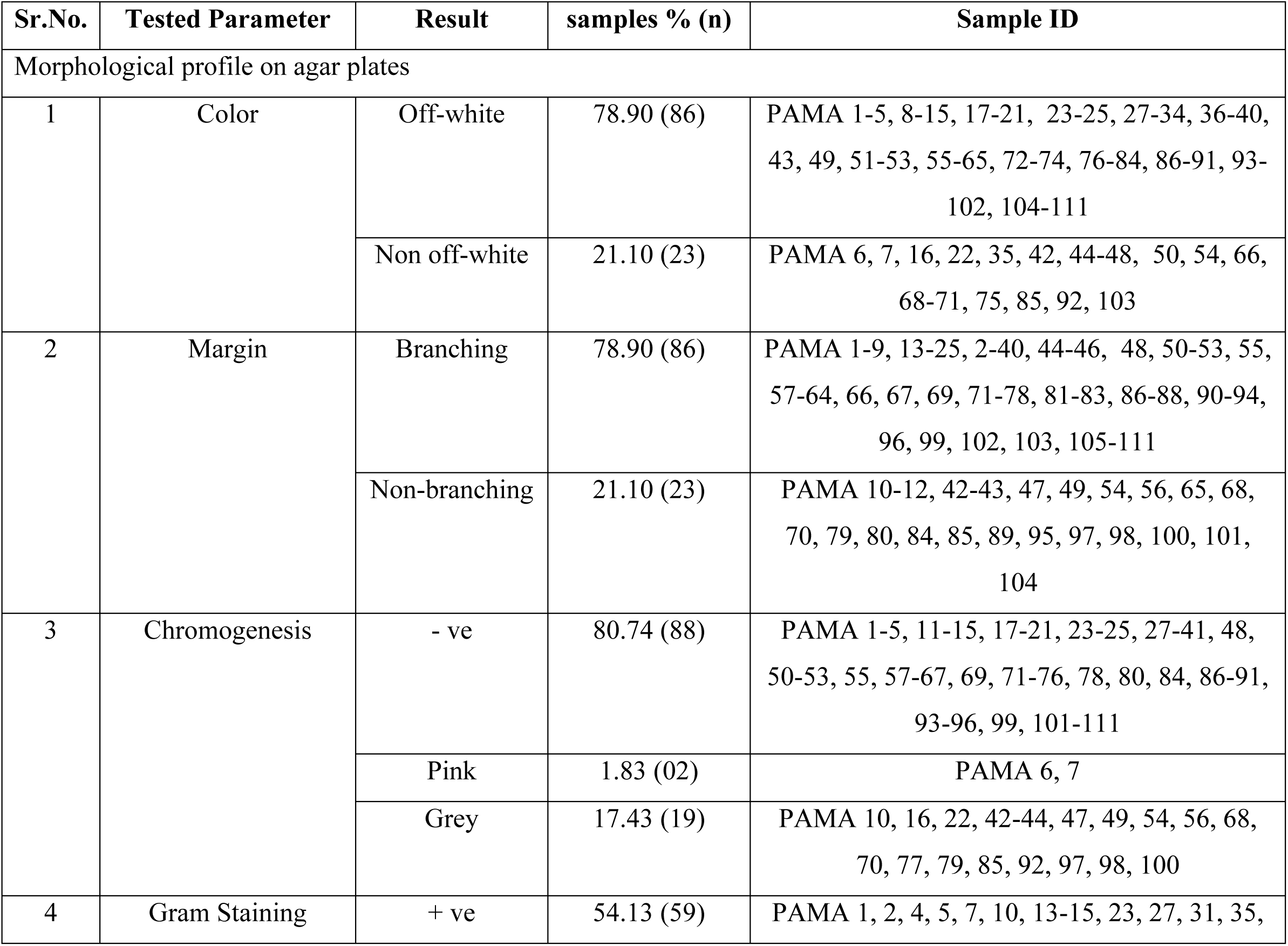
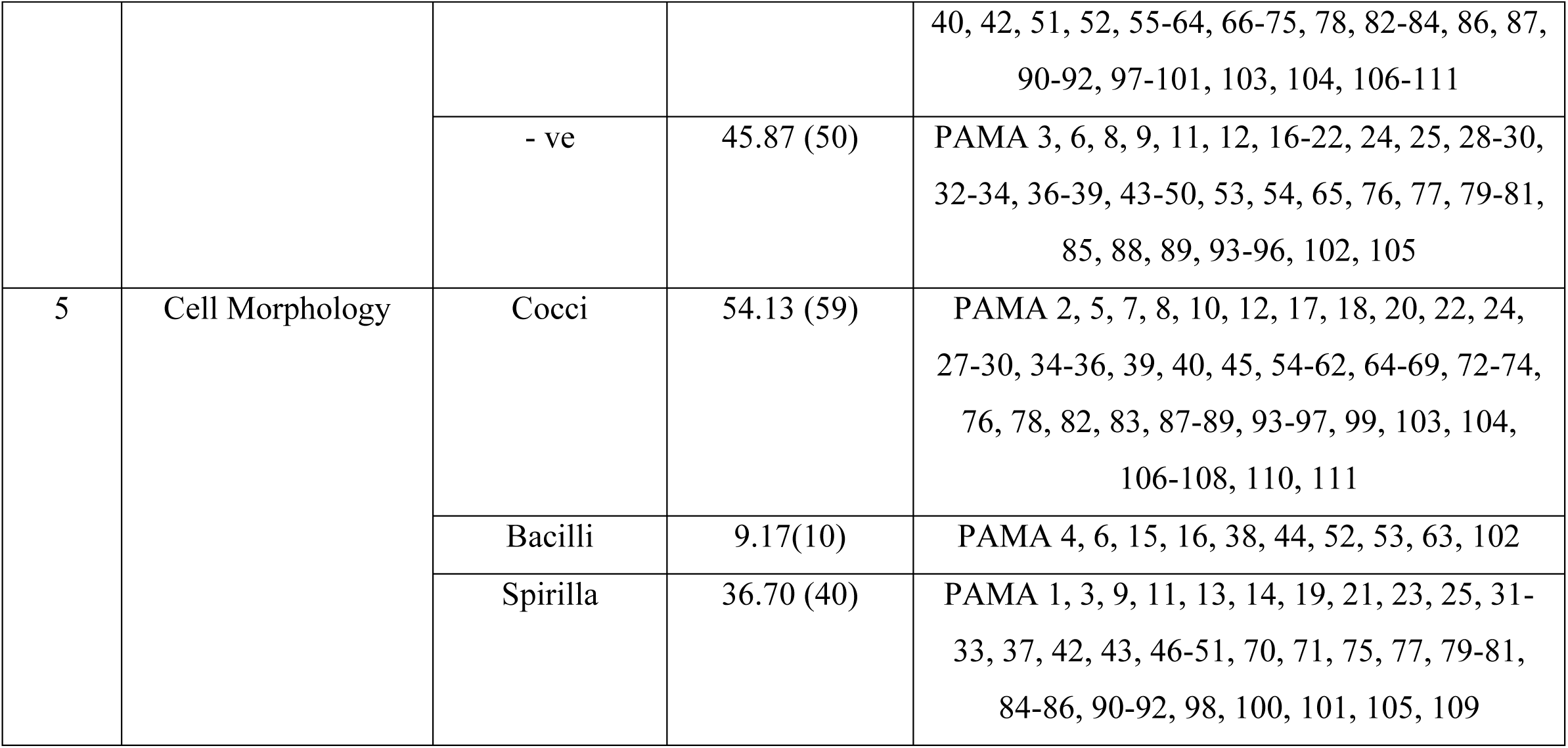
Morphological profile of bacteria (PAMA Series) isolated from rhizosphere of *Panicum antidotale*.

| Sr.No. | Tested Parameter | Result | samples % (n) | Sample ID |
| --- | --- | --- | --- | --- |
| Morphological profile on agar plates |  |  |  |  |
| 1 | Color | Off-white | 78.90 (86) | PAMA 1-5, 8-15, 17-21, 23-25, 27-34, 36-40, 43, 49, 51-53, 55-65, 72-74, 76-84, 86-91, 93-102, 104-111 |
|  |  | Non off-white | 21.10 (23) | PAMA 6, 7, 16, 22, 35, 42, 44-48, 50, 54, 66, 68-71, 75, 85, 92, 103 |
| 2 | Margin | Branching | 78.90 (86) | PAMA 1-9, 13-25, 2-40, 44-46, 48, 50-53, 55, 57-64, 66, 67, 69, 71-78, 81-83, 86-88, 90-94, 96, 99, 102, 103, 105-111 |
|  |  | Non-branching | 21.10 (23) | PAMA 10-12, 42-43, 47, 49, 54, 56, 65, 68, 70, 79, 80, 84, 85, 89, 95, 97, 98, 100, 101, 104 |
| 3 | Chromogenesis | - ve | 80.74 (88) | PAMA 1-5, 11-15, 17-21, 23-25, 27-41, 48, 50-53, 55, 57-67, 69, 71-76, 78, 80, 84, 86-91, 93-96, 99, 101-111 |
|  |  | Pink | 1.83 (02) | PAMA 6, 7 |
|  |  | Grey | 17.43 (19) | PAMA 10, 16, 22, 42-44, 47, 49, 54, 56, 68, 70, 77, 79, 85, 92, 97, 98, 100 |
| 4 | Gram Staining | + ve | 54.13 (59) | PAMA 1, 2, 4, 5, 7, 10, 13-15, 23, 27, 31, 35, |
|  |  |  |  | 40, 42, 51, 52, 55-64, 66-75, 78, 82-84, 86, 87, 90-92, 97-101, 103, 104, 106-111 |
|  |  | - ve | 45.87 (50) | PAMA 3, 6, 8, 9, 11, 12, 16-22, 24, 25, 28-30, 32-34, 36-39, 43-50, 53, 54, 65, 76, 77, 79-81, 85, 88, 89, 93-96, 102, 105 |
| 5 | Cell Morphology | Cocci | 54.13 (59) | PAMA 2, 5, 7, 8, 10, 12, 17, 18, 20, 22, 24, 27-30, 34-36, 39, 40, 45, 54-62, 64-69, 72-74, 76, 78, 82, 83, 87-89, 93-97, 99, 103, 104, 106-108, 110, 111 |
|  |  | Bacilli | 9.17(10) | PAMA 4, 6, 15, 16, 38, 44, 52, 53, 63, 102 |
|  |  | Spirilla | 36.70 (40) | PAMA 1, 3, 9, 11, 13, 14, 19, 21, 23, 25, 31-33, 37, 42, 43, 46-51, 70, 71, 75, 77, 79-81, 84-86, 90-92, 98, 100, 101, 105, 109 |

### Biochemical characteristics

Biochemical profiling of 108 bacterial isolates revealed distinct patterns (Table 2). Amylase production was positive in 37.04% (40 isolates) and negative in 55.56% (60 isolates). Acid production was observed in 25.92% (28 isolates), while urease production was low, with only 8.33% (9 isolates) testing positive. Gelatinase production was absent in all isolates (0% positive). Motility was detected in 11.11% (12 isolates). The Methyl Red (MR) test was positive in 41.67% (45 isolates). Voges-Proskauer (VP) reaction and H₂S production were not detected in any isolate (0% positive for both). Catalase production was universally present (100% positive). All isolates fermented glucose, lactose, and fructose (100% each).

**Table 2.** Biochemical characteristics of bacteria (PAMA Series) isolated from the rhizosphere of *Panicum antidotale*.

| Sr.No. | Tested Parameter | Result | samples<br>% (n) | Sample ID |
| --- | --- | --- | --- | --- |
| 1 | Amylase production | + ve | 44.95 (49) | PAMA 1-9, 11-15, 23, 27-32, 51-53, 56-61, 72-75, 80-85, 93-95, 101-106 |
|  |  | - ve | 55.05 (60) | PAMA 10, 16-22, 24, 25, 33-40, 42-50, 54, 55, 62-71,<br>76-79, 86-92, 96-100, 107-111 |
| 2 | Acid production | + ve | 23.85 (26) | PAMA 10, 16-19, 24-25, 33-35, 54, 55, 62, 63, 76-79,<br>86, 96-100, 107, 108 |
|  |  | - ve | 76.15 (83) | PAMA 1-9, 11-15, 20-23, 27-32, 36-40, 42-53, 56-61,<br>64-75, 80-85, 87-95, 101-106, 109-111 |
| 3 | Urease production | + ve | 12.84 (14) | PAMA 1, 3, 4, 6, 9, 10, 23, 36, 54, 72, 73, 75, 88, 95 |
|  |  | -ve | 87.16 (95) | PAMA 2, 5, 7, 8, 11-22, 24-35, 37-40, 42-53, 55-71,<br>74, 76-87, 89-94, 96-111 |
| 4 | Gelatinase production | +ve | 4.59 (5) | PAMA 20, 36, 39, 65, 91 |
|  |  | -ve | 95.41 (104) | PAMA 1-19, 21-25, 27-35, 37, 38, 40, 42-64, 66-90,<br>92-111 |
| 5 | Protease production | + ve | 15.60 (17) | PAMA 20, 21, 36-40, 64, 65, 87-91, 109-111 |
|  |  | - ve | 84.40 (92) | PAMA 1-19, 22-25, 27-35, 42-63, 66-86, 92-108 |
| 6 | Indol production | + ve | 11.93 (13) | PAMA 22, 42-48, 66-69, 92 |
|  |  | - ve | 88.07 (96) | PAMA 1-21, 23-25, 27-40, 49-65, 70-91, 93-111 |
| 7 | H<sub>2</sub>S production | + ve | 3.67 (4) | PAMA 49, 50, 70, 71 |
|  |  | - ve | 96.33 (105) | PAMA 1-25, 27-40, 42-48, 51-69, 72-111 |
| 8 | Glucose fermentation | + ve | 20.18 (22) | PAMA 1, 2, 10, 15, 17, 19, 24, 27, 30, 32, 33, 35, 52,<br>54, 55, 76, 79, 81, 82, 98, 109, 110 |
|  |  | -ve | 79.82 (87) | PAMA 3-9, 11-14, 16, 18, 20-23, 25, 28, 29, 31, 34,<br>36-51, 53, 56-75, 77, 78, 80, 83-97, 99-108, 111 |
| 9 | Lactose fermentation | + ve | 57.80 (63) | PAMA 2, 5, 7, 9-25, 27-34, 38, 51, 52, 54-56, 59, 61,<br>65, 67, 69, 71, 76-81, 83-86, 88, 89, 96-100, 104, 105,<br>107-109, 111 |
|  |  | -ve | 42.20 (46) | PAMA 1, 3, 4, 6, 8, 35-37, 39, 40, 42-50, 53, 57, 58,<br>60, 62-64, 66, 68, 70, 72-75, 82, 87, 90-95, 101-103,<br>106, 110 |
| 10 | Fructose fermentation | + ve | 17.43 (19) | PAMA 7, 10, 14, 18, 24, 25, 29, 36, 40, 51, 55, 76-78,<br>86, 96, 98, 102, 111 |
|  |  | -ve | 82.57 (90) | PAMA 1-6, 8, 9, 11-13, 15-17, 19-23, 27, 28, 30-35,<br>37-39, 42-50, 52-54, 56-75, 79-85, 87-95, 97, 99-101, |
|  |  |  |  | 103-110 |
| 11 | Ulrich milk test | + ve | 77.98(85) | PAMA 1-5, 7, 9-21, 23-25, 27-40, 51-65, 76-91, 93-102, 104-111 |
|  |  | -ve | 22.02 (24) | PAMA 6, 8, 22, 26, 42-50, 66-75, 92, 103 |

### Molecular characterization

Molecular characterization was performed using three techniques: 16S rRNA profiling, RFLP, and DNA sequence analysis. Partial 16S rRNA gene sequences (∼1100 bp) were analyzed using BLAST to identify bacterial species. The sequences matched known bacteria with varying similarity percentages. Interestingly, the phylogenetic trees generated from RFLP patterns and from the DNA sequences did not perfectly match, although RFLP is a recognized method for species-level differentiation [15].

BLAST, a widely used algorithm for nucleotide and protein sequence analysis [16], identified three bacterial groups in the samples (Among the 20 bacterial strains analyzed, 6 were identified as *Bacillus subtilis*, 9 as *Bacillus licheniformis*, 3 as *Brevibacillus borstelensis*, and 2 as *Paenibacillus dendritiformis*. Multiple sequence alignment was performed using Geneious [17]. Truncated sequences were removed, and longer sequences were shortened to ensure uniform length.

In the phylogenetic tree constructed using 16S rRNA gene sequences, three main clades were observed: *Bacillus* spp., *Paenibacillus dendritiformis*, and *Brevibacillus borstelensis*. The *Bacillus* clade contained two species: *Bacillus subtilis* and *Bacillus licheniformis*. Bootstrap values and clustering patterns indicated a close evolutionary relationship between these two *Bacillus* species. Phylogenetic analysis also revealed that *Paenibacillus dendritiformis* was more closely related to *Bacillus* spp. than to *Brevibacillus borstelensis*. The overall evolutionary distance (d) and standard error (S.E.) were 0.136 and 0.006, respectively, with standard error estimates obtained through bootstrap analysis.

The Neighbor-Joining method [18] was used to infer evolutionary history, and a bootstrap consensus tree was generated from 100 replicates. Branches reproduced in fewer than 50% of bootstrap replicates were collapsed. The percentage of replicate trees in which the associated taxa clustered together is shown next to the branches [19]. A total of 923 positions were included in the final dataset, and evolutionary analyses were performed in MEGA7 [20].

To understand the evolutionary relationships of the bacterial isolates, a phylogenetic tree was constructed using the Neighbor-Joining method. This tree was based on aligned 16S rRNA sequences from both our isolates and their closest relatives retrieved from the NCBI GenBank database. This approach was essential to examine their evolutionary relationships and clade patterns.

Based on the Neighbor-Joining phylogenetic tree generated from the aligned 16S rRNA sequences, BLAST analysis identified three major bacterial groups in our samples (Table 3). Among the 20 bacterial strains analyzed, 6 were identified as *Bacillus subtilis*, 9 as *Bacillus licheniformis*, 3 as *Brevibacillus borstelensis*, and 2 as *Paenibacillus dendritiformis*. Multiple sequence alignment (Figure 1) was performed using Geneious [17]. Truncated sequences were removed, and longer sequences were trimmed to ensure uniform length across the alignment.

**Figure 1:**
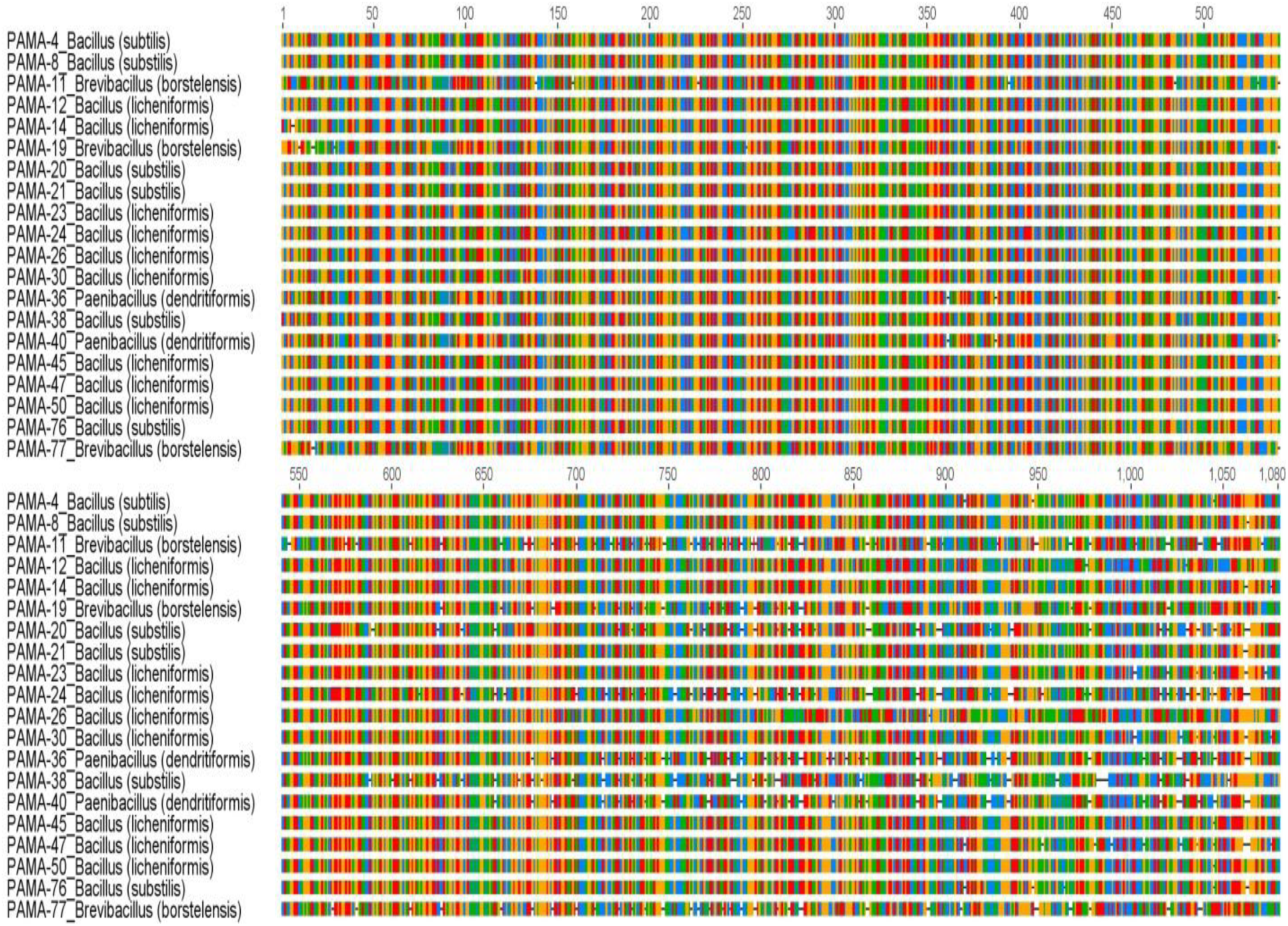
Multiple sequence alignment of 16S rRNA genes of bacteria isolated from rhizosphere of *Panicum antidotale* (PAMA Series). Similar color bars are showing similar sequences. Consensus sequence is also shown in the alignment.

**Table 3:**
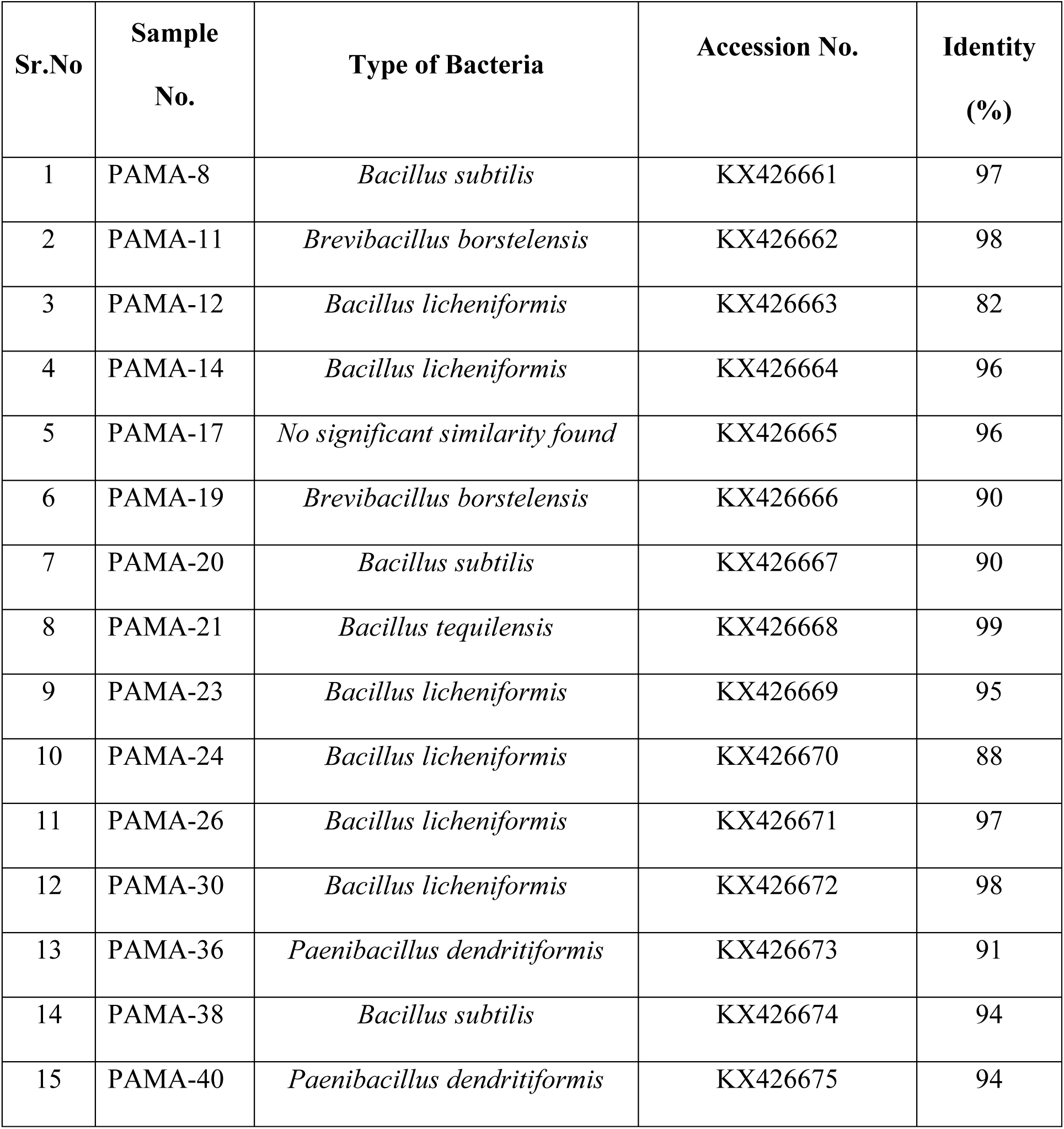
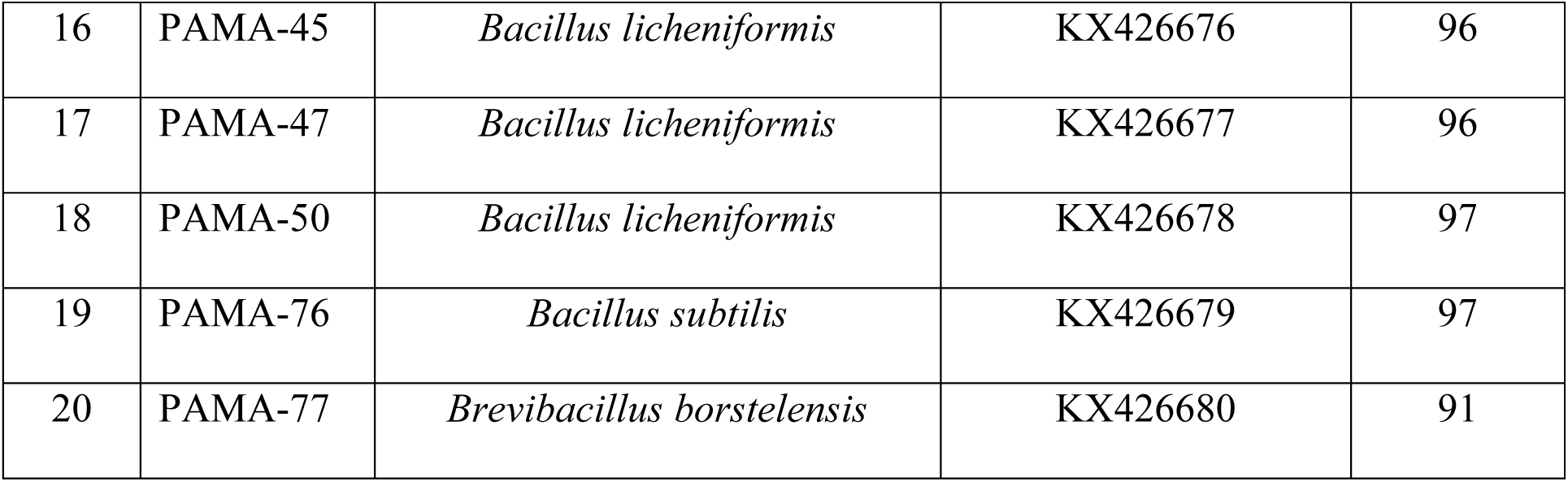
Nearest relatives of selected bacterial isolates (PAMA Series) and their accession numbers, collected from the rhizosphere of *Panicum antidotale*.

In the constructed phylogenetic tree, three main clades were observed (Figure 2): *Bacillus* spp., *Paenibacillus dendritiformis*, and *Brevibacillus borstelensis*. The *Bacillus* clade consisted of two species, *Bacillus subtilis* and *Bacillus licheniformis*. Bootstrap values and branch positions indicated a close evolutionary relationship between these two *Bacillus* species. Phylogenetic analysis further revealed that *Paenibacillus dendritiformis* was more closely related to *Bacillus* spp. than to *Brevibacillus borstelensis*.

**Figure 2.**
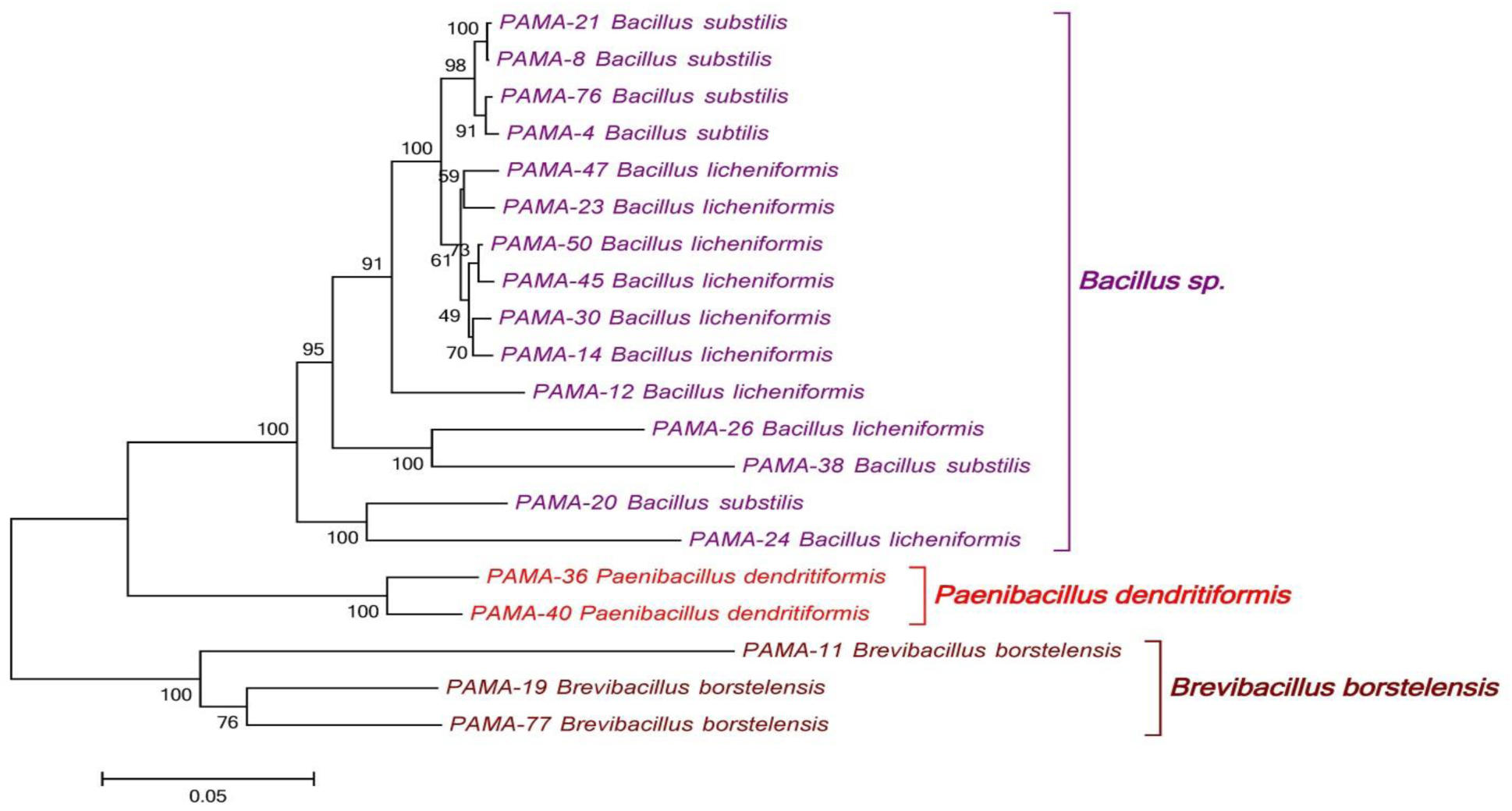
Molecular phylogenetic analysis of 16S rRNA genes of bacteria isolated from rhizosphere of *Panicum antidotale* (PAMA Series)

The overall evolutionary distance (d) and standard error (S.E.) were found to be 0.136 and 0.006, respectively. Standard error estimates were obtained through a bootstrap procedure. The Neighbor-Joining method [18] was used to infer the evolutionary history of the analyzed taxa, represented by a bootstrap consensus tree generated from 100 replicates. Branches reproduced in fewer than 50% of bootstrap replicates were collapsed. The percentage of replicate trees in which the associated taxa clustered together during the bootstrap test is shown next to the branches [19]. The final dataset contained a total of 923 positions, and evolutionary analyses were conducted using MEGA7 [20].

### RFLP and phylogenetic analysis

PCR amplicons of 16S rRNA genes were digested using five restriction endonucleases (i.e., RsaI, HinfI, TaqI, HpaI, and HhaI), and the resulting restriction fragments were resolved on agarose gel (Figure 3). A dendrogram was constructed based on the RFLP data, which revealed three main clades: A, B, and C.

**Figure 3.**
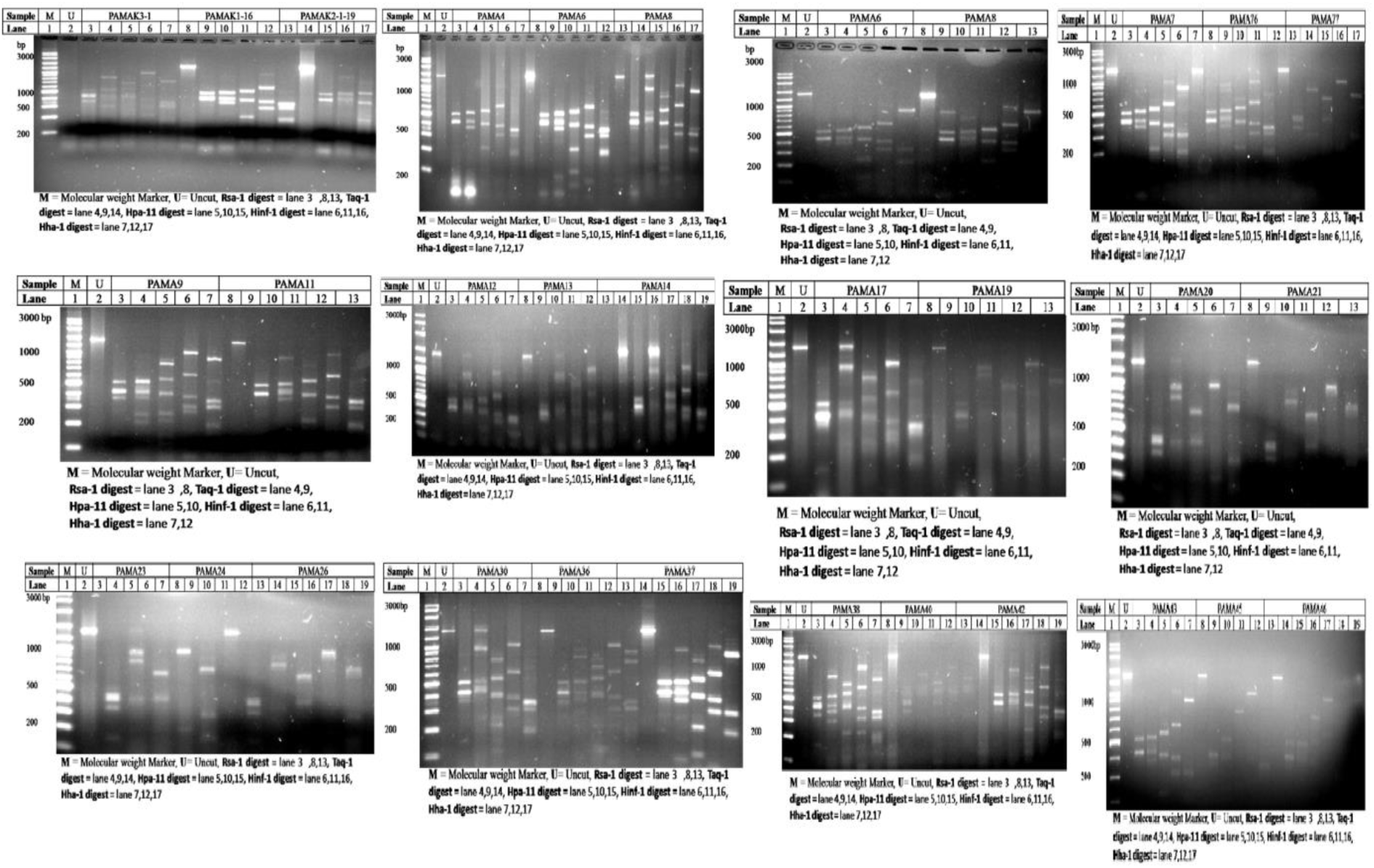
RFLP analysis of some bacterial isolates (PAMA 99, 107, 109) from the rhizosphereof *Panicum antidotale*

Clade A and Clade B, each contained a single isolate, PAMA 21 and PAMA 20, respectively. The remaining 107 isolates were grouped into Clade C (Figure 4). Clade C was further divided into two sub-clades: c1 and c2. Sub-clade c1 was composed of two sub-sub-clades: c1.1 and c1.2. The major sub-sub-clade, c1.1, consisted of 102 isolates, while c1.2 comprised five isolates (i.e., PAMA 6-MPhil, 10-MPhil, K1-16, K2-1-16, K2-1-19, and K2-1-26).

**Figure 4.**
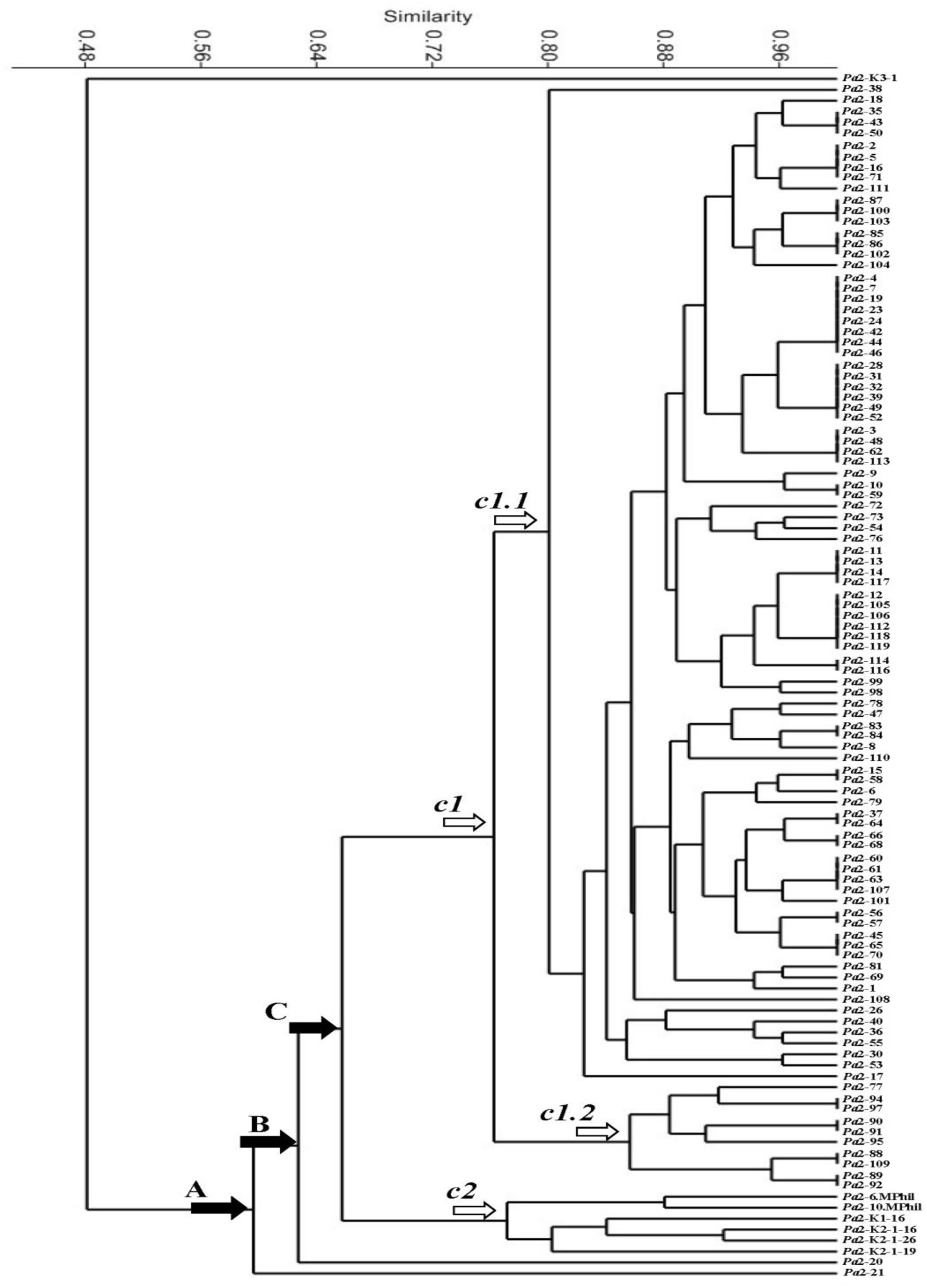
Dendrogram constructed on the basis of RFLP analysis of 16S rRNA gene of bacteria (PAMA series) isolated from rhizosphere of *Panicum antidotale*

Clades A, B, and C were found to be equally distant yet closely related, sharing a common ancestor. Among the 109 isolates, one isolate (PAMA K3-1) appeared to have evolved from a unique ancestor and did not cluster within Clades A, B, or C.

The presence of multiple isolates with identical RFLP patterns in sub-sub-clade c1.1 suggested a high level of genetic similarity. To resolve the ambiguity in the identification of these isolates, the study was further extended to include DNA sequence analysis of 20 isolates, randomly selected to represent all observed RFLP patterns (Figure 5).

**Figure 5.**
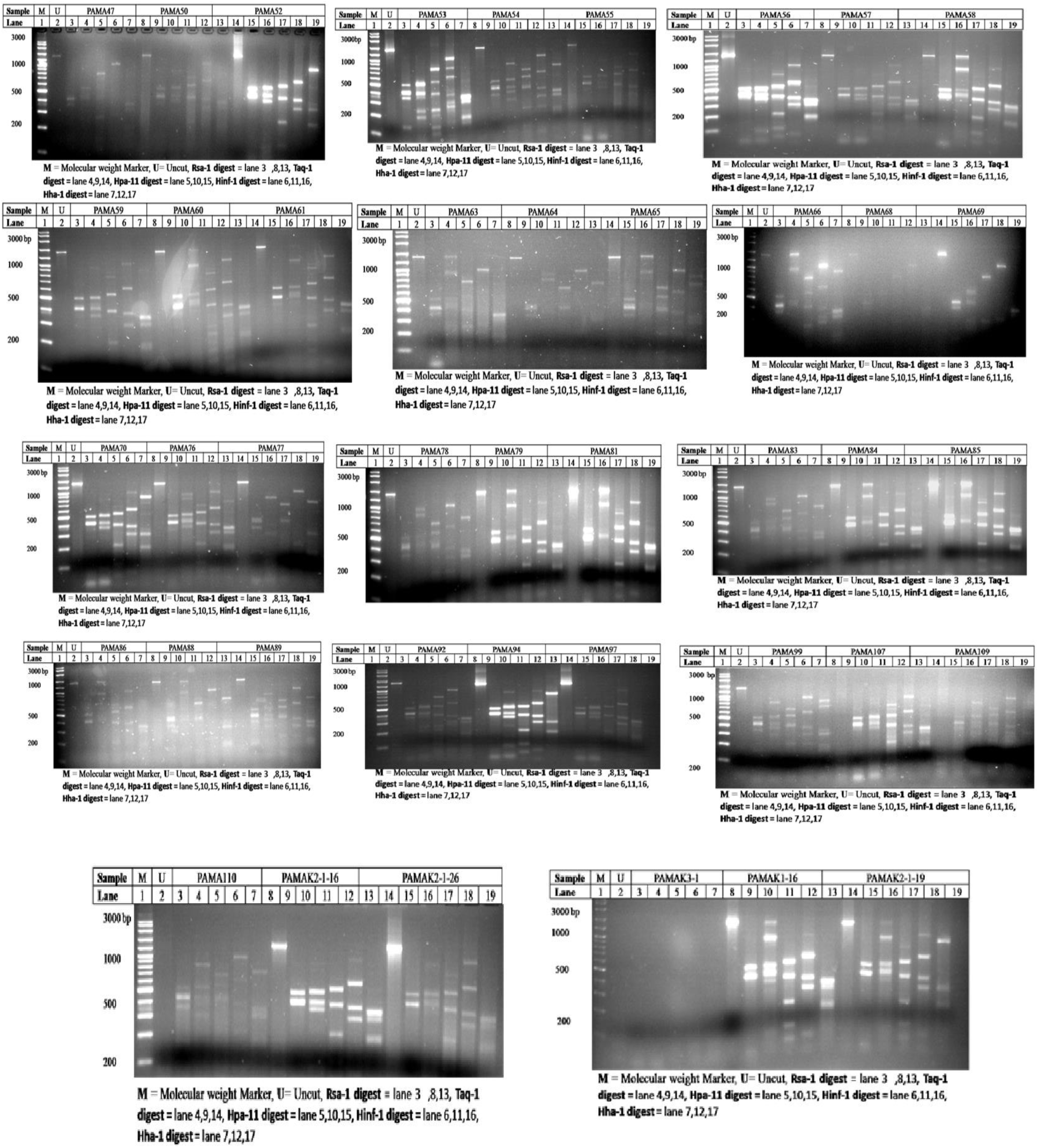
RFLP analysis of some bacterial isolates (PAMA 99, 107, 109) from the rhizosphere of *Panicum antidotale*

## Discussion

In the present study, *Bacillus* species were identified as the predominant members of the microbial community associated with the rhizosphere of *Panicum antidotale* in the Cholistan Desert. Plant growth-promoting bacteria (PGPB) are known to enhance the growth of desert plants by improving soil stabilization, reducing erosion, and exerting beneficial effects on plant metabolism. These effects may occur directly (e.g., production of phytohormones, nutrient solubilization) or indirectly by suppressing phytopathogens such as bacteria, fungi, viruses, and nematodes [21, 22]. Despite the ecological importance of desert ecosystems, the rhizospheres of desert plants remain poorly studied globally [23]. To our knowledge, this is the first report on the isolation and characterization of culturable bacteria from the rhizosphere of *P. antidotale* in Pakistan’s Cholistan Desert.

The bacterial isolates identified from the rhizosphere of *P. antidotale* included *Brevibacillus borstelensis*, *Bacillus subtilis*, *Bacillus licheniformis*, *Paenibacillus dendritiformis*, and in some samples, *Pseudomonas fluorescens*. Among these, *Bacillus* species were dominant, with *B. subtilis* and *B. licheniformis* being the most abundant, corroborating their well-documented role as aerobic, endospore-forming PGPB. These species are known to produce valuable extracellular enzymes such as proteases, amylases, laccases, and lipases, and they can degrade complex carbohydrates including cellulose, xylulose, and oligosaccharides like arabinogalactan, stachyose, and raffinose [24, 25].

Thermophilic *Bacillus* and *Brevibacillus* strains have also demonstrated antimicrobial activity against both Gram-positive bacteria (*Micrococcus luteus*, *Staphylococcus aureus*) and Gram-negative bacteria (*Pseudomonas aeruginosa*, *Escherichia coli*, *Klebsiella pneumoniae*) [26]. *B. subtilis* produces secondary metabolites, including amocoumacin-A-like compounds, with antimicrobial potential against fungi (*Candida albicans*, *Ustilago maydis*, *Cryptococcus neoformans*) and pathogenic bacteria [27]. Both *Brevibacillus borstelensis* and *B. licheniformis* have been reported to induce systemic resistance in plants and inhibit phytopathogens [28, 29], although *B. licheniformis* has occasionally been associated with opportunistic infections in humans [30].

Interestingly, *Brevibacillus* species also exhibit bioremediation potential, being capable of degrading toxic chemicals such as toluidine blue dye and the fungicide carbendazim [31–33]. *Pseudomonas fluorescens*, another isolate from our samples, is a well-known non-pathogenic saprophyte commonly found in soil, water, and plant surfaces [34]. Many *Pseudomonas* strains act as biocontrol agents by producing siderophores, hydrogen cyanide, and antibiotics that suppress fungal pathogens and promote plant health [35–37].

*Paenibacillus dendritiformis* is an endospore-forming facultative anaerobe formerly classified under *Bacillus*. Members of this genus are widespread in rhizospheres, soil, and water [38, 39], and they produce exopolysaccharides, proteases, and polysaccharide-degrading enzymes used in wastewater treatment and industrial processes [40]. Several *Paenibacillus* species also synthesize antimicrobial compounds with activity against plant and soil pathogens [41, 42].

Other detected genera, including *Stenotrophomonas* spp., are known for their biocontrol and plant growth-promoting properties [43, 44]. *Stenotrophomonas rhizophila*, for instance, protects plants against fungal diseases and produces osmoprotectants such as glucosylglycerol and trehalose, enhancing plant stress tolerance [45, 46]. Their ability to degrade xenobiotic compounds also positions them as promising candidates for bioremediation [47].

Conversely, *Pseudomonas marginalis*, detected in some samples, is a fluorescent bacterium known to degrade pectin and cause soft rot in stored vegetables and necrosis in plants [48–50]. Its presence suggests the coexistence of plant-pathogenic bacteria within the rhizosphere of Cholistan desert plants, highlighting the dynamic nature of this microbial community.

Overall, the presence of beneficial bacterial taxa—particularly *Bacillus* spp., *Brevibacillus*, *Paenibacillus*, and *Pseudomonas fluorescens*—indicates a potential natural defense system in the rhizosphere of *P. antidotale*, contributing to plant growth promotion and protection under arid conditions. However, the isolation-based approach used in this study may have captured only a subset of the true microbial diversity, as many rhizosphere microorganisms remain unculturable under standard laboratory conditions. Future studies integrating metagenomic and functional assays will be essential to unravel the complete microbiome and its role in desert plant adaptation and ecosystem resilience.

## Declarations

## Authors’ Contributions

F.-u.-H. Nasim served as the principal investigator (PI) and HEC grant recipient. M. Aslam, M. Afzal, and R. Batool collaborated on sample identification, collection, and analyses. M. Aslam and R. Batool performed the bench work. S. Ejaz, F.-u.-H. Nasim, T. Rehman, M. Aslam, and M. Afzal contributed to data analyses and manuscript writing. M. Aslam prepared the dendrograms and conducted the related analyses. T. Rehman served as the corresponding author and contributed to manuscript revision and final approval.

## Acknowledgements

This study was supported by a grant from the Higher Education Commission (HEC), Pakistan, awarded to F.-u.-H. Nasim, which is gratefully acknowledged.

## Availability of Data and Materials

Ribosomal DNA sequences have been submitted to GenBank under accession numbers provided in Table 3.

## Conflict of Interest

The authors declare that they have no financial or non-financial competing interests.

